# Fe-TAMLs as a new class of small molecule peroxidase probes for correlated light and electron microscopy

**DOI:** 10.1101/2023.08.25.554352

**Authors:** Stephen R. Adams, Mason R. Mackey, Ranjan Ramachandra, Thomas J. Deerinck, Guillaume A. Castillon, Sebastien Phan, Junru Hu, Daniela Boassa, John T. Ngo, Mark H. Ellisman

## Abstract

We introduce Fe-TAML, a small molecule-based peroxidase as a versatile new member of the correlated fluorescence and electron microscopy toolkit. The utility of the probe is demonstrated by high resolution imaging of newly synthesized DNA (through biorthogonal labeling), genetically tagged proteins (using HaloTag), and untagged endogenous proteins (via immunostaining). EM visualization in these applications is facilitated by exploiting Fe-TAML’s catalytic activity for the deposition of localized osmiophilic precipitates based on polymerized 3,3’-diaminobenzidine. Optimized conditions for synthesizing and implementing Fe-TAML based probes are also described. Overall, Fe-TAML is a new chemical biology tool that can be used to visualize diverse biomolecular species along nanometer and micron scales within cells.

## INTRODUCTION

Probes and tags for visualizing biomolecules by light and electron microscopy are powerful tools for dissecting the molecular organization of cell[1,2]. Over recent years, molecular-engineering based strategies been used to develop new labels to track specific proteins via EM, including small taggable peptide sequences or proteins[3–6], fluorescent proteins capable of generating EM-visible reaction products [7] as well as enzyme-based tags such as encodable peroxidases HRP and APEX2 [8–10]. Despite these developments, a need for new probes remains - especially for those that can be used to track diverse biomolecules, including endogenous and non-protein based species. In particular, versatile labels of small molecular dimensions (for maximal resolving precision), as well as those that can be combined with extant probes (for ’multicolor’ analyses), are especially desirable and missing components of the existing toolset.

Biomolecule localization via EM has traditionally been accomplished using antibodies, with EM contrast being introduced via one of two modes: with nanoparticulate colloidal gold, or through the in-situ polymerization of 3,3’-diaminobenzidine (DAB) using antibodies conjugated with HRP (horseradish peroxidase). However, both of these labels carry serious limitations due to the size of the resulting conjugate. For a whole IgG conjugated to HRP it is ∼190 kDa and for IgG-colloidal gold conjugates, gold particles larger than 3 nm. Such bulky tagging schemes can be problematic by limiting labeling efficiency through poor penetration into fixed samples that retain the cellular ultrastructure.

To overcome the disadvantages associated with large labels such as HRP, we reasoned that a small molecule-based catalyst capable of facilitating a similar *in situ* polymerization of DAB would provide a new and versatile strategy to labeled diverse biomolecular species for imaging by EM. To develop such a probe, we thus turned to Fe-TAMLs (for *Fe(III) tetra-amido macrocyclic ligands*), a family of small synthetic peroxidase mimics originally designed to serve as efficient catalysts for oxidative reactions. Given the ’green chemistry’ interest of using Fe-TAMLs for destroying environmental pollutants, including aquatic water sources, various efforts have been made to optimize its activity and stability in aqueous media [11,12]. Fe-TAML has at its core an iron atom surrounded by four nitrogen atoms, which in turn are bound by a ring of carbon atoms (Figure 1A). Water molecules loosely associate with the vertical pole of the iron atom, but in the presence of hydrogen peroxide, the water is displaced and the complex acts as a catalyst that triggers oxidation reactions with other compounds in solution.

**Figure 1.**
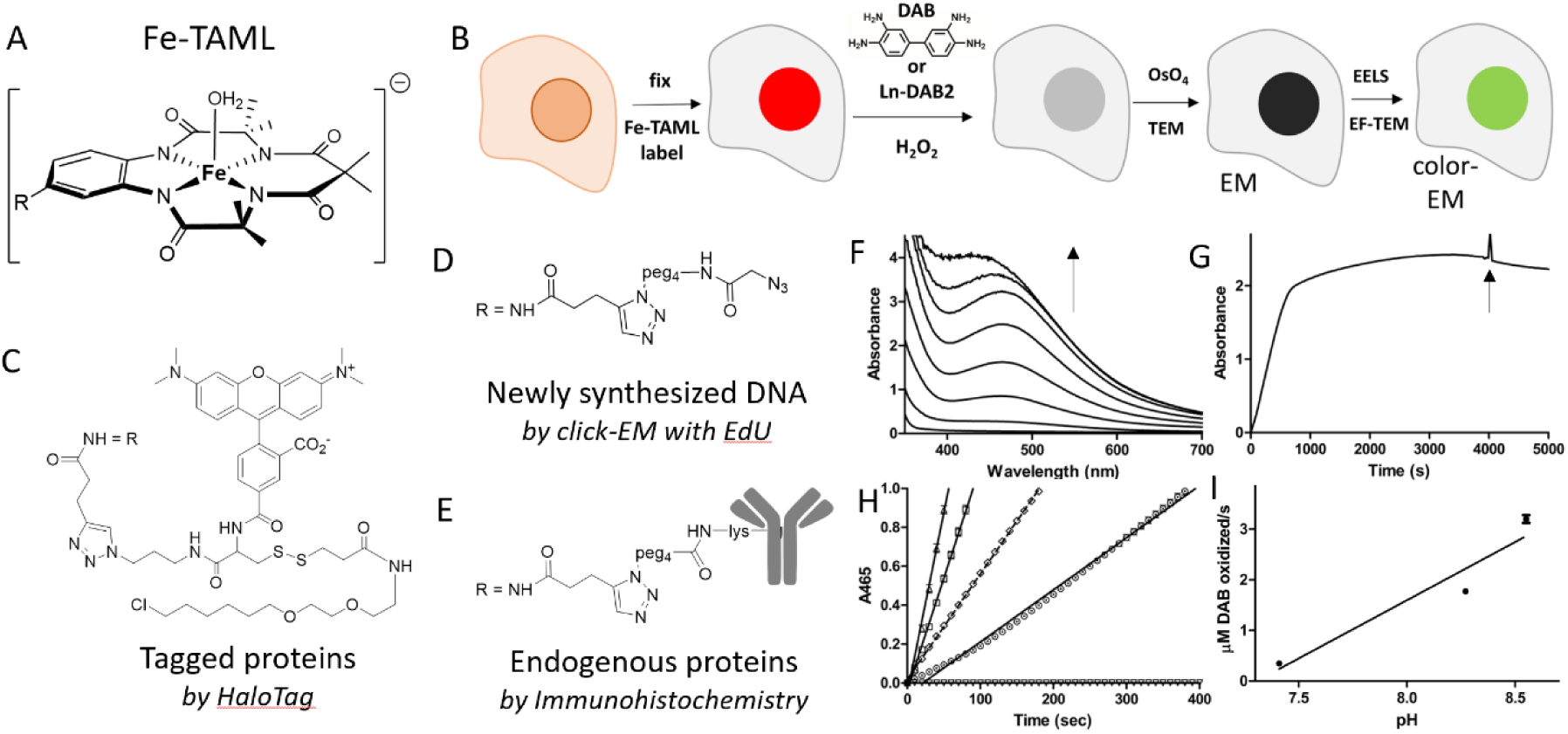
Application of Fe-TAML derivatives to correlated light and electron microscopy. A) Chemical structure of Fe-TAML B* (R is -H) and Fe-TAML alkyne (R is -NHCOCH_2_CH≡CH). B) Scheme of cell labeling procedure with Fe-TAML conjugates for conventional EM or color EM. C-E) Chemical structures of Fe-TAML derivatives for labeling transfected proteins, DNA, and endogenous proteins respectively. F) Consecutive absorbance spectra upon oxidation of 0.5 mM DAB by 10 mM H_2_O_2_ catalyzed by 5 μM Fe-TAML B* at pH 7.4; 1, 3, 5, 8, 17 and 32 min after addition of Fe-TAML to DAB and H_2_O_2_. G) Time course at 465 nm of DAB oxidation as in F but with pathlength of 0.5 cm. An additional 5 μM Fe-TAML B* and 10 mM H_2_O_2_ were added as indicated by arrow. H) Fitted initial rates from progress curves of 0.5 mM DAB oxidation by 40 mM H_2_O_2_ with 0.5 μM Fe-TAML at pH 7.41 (circles, solid line), 8.27 (squares, solid line) and 8.55 (triangle, solid line). The reaction without Fe-TAML at pH 8.55 is essentially baseline (inverted triangles) and for comparison, the reaction catalyzed by 1 nM horse radish peroxidase with 1 mM H_2_O_2_ (circles, dotted line) at pH 7.4. I) pH dependance of DAB oxidation catalyzed by Fe-TAML B*.

Reasoning that Fe-TAML could be adapted as a small molecule catalyst for EM (by catalyzing DAB oxidation at sites where the probe is targeted), we thus tested the catalyst in solution and cell-based reactions and identified optimal conditions for its use in polymerizing DAB. We show that Fe-TAML can be utilized in EM labeling-based application via procedures that are mechanistically analogous to that which are used with HRP, but in ways that can be extended to diverse molecular species (including viral proteins and newly replicated DNA). Design strategies and synthetic procedures for preparing multimodal Fe-TAML probes are also described, including a click-reactive peroxidase mimic, a trifunctional and fluorescent HaloTag ligand, as well as secondary catalytic and fluorescent antibody conjugates. Overall, the main advantages of Fe-TAMLs are that they are very small (M.W. ∼500), about 1% the size of HRP by mass, are water soluble, capable of large turnover numbers, are extremely stable under a wide range of pH and ionic conditions and are amenable to conjugation for targeted subcellular labeling. Furthermore, since they are completely synthetic, chemical modification can be used to generate powerful, selective catalytic probes that can be targeted to multiple cellular labeling targets of interest.

Herein, we provide validation of Fe-TAML as a versatile labeling probe for marking specified biomolecules for visualization by CLEM (Figure 1B). We demonstrate utility of the probe in several ways, including use of a bioorthogonally functionalized Fe-TAML in click-chemistry mediated imaging of metabolically tagged DNA as well as use of a haloalkane functionalized probe to reveal HaloTag-tagged protein constructs (Figure 1C-D). Further utility of the system is demonstrated through the EM detection of endogenously expressed untagged proteins, as revealed using antibodies conjugated with the Fe TAML (Figure 1E).

## RESULTS AND DISCUSSION

### Fe-TAML catalyzed oxidative precipitation of DAB with H_2_O_2_

A related biuret Fe-TAML [13] has been reported to catalyze the oxidation by H_2_O_2_ of the related chromophoric but non-precipitating benzidine derivative, TMB (3,5,3’,5’-tetramethylbenzidine), for use in glucose detection[14] and for in gel-detection of Fe-TAML protein conjugates[15]. We therefore tested the ability and kinetics of the prototypical Fe-TAML B* (Figure 1A, R=H) to catalyze DAB oxidation and precipitation by H_2_O_2_ in a pH 7.4 cacodylate buffer conventionally used for high quality fixed TEM cell samples. With micromolar Fe-TAML, a 2.5 mM solution of DAB rapidly turned brown on addition of 10 mM H_2_O_2,_ and a dense precipitate, consistent with the anticipated reaction product, was formed. To determine the catalytic efficacy, we carried out a previously reported colorimetric peroxidase assay in which DAB is reacted in solution containing 0.1% (w/v) gelatin as an oxidation product solubilizer. Using this solution we monitored the reaction progress until the DAB substrate was fully consumed[16]. Addition of micromolar amounts of Fe-TAML to millimolar H_2_O_2_ and 0.5 mM DAB gave an increase in absorbance at 465 nm, characteristic of the oxidation product of DAB[16] (Figure 1F). After 65 minutes, addition of fresh Fe-TAML and H_2_O_2_ did not result in any further increases in absorbance at 465 nm, indicating complete consumption of the starting DAB (Figure 1G). This result verified the ability for Fe-TAML to facilitate high turnover of DAB substrates. In the absence of Fe-TAML, the solution remained colorless, verifying the catalytic activity of Fe-TAML in DAB oxidation. The reaction displayed Michaelis-Menten kinetics from the initial linear rates (Figure 1H) giving *K*_M_ values of 110 mM for H_2_O_2_ and 87 μM for DAB with a decrease in rates at higher than 1 mM DAB (Figure S3).

Previous work has shown that Fe-TAML catalyzed conversion rates can vary according to pH with a 5-to 6-fold increase in rate of Fe-TAML catalyzed oxidation of the azo dye Orange II at pH 9 compared to pH 7[13]. Using such non-physiological pH are possible options for generating DAB precipitation in fixed cells that have been treated with the high concentrations of aldehydes (paraformaldehyde and/or glutaraldehyde) that preserve optimal cellular ultrastructure for TEM. To determine whether elevated pH would facilitate increased DAB precipitation by Fe-TAML, we tested oxidation rates in pH 8.23 and 8.55 bicine buffered solutions (Figure 1H). As expected, we observed rate enhancements of 5.1- and 9.2-fold, respectively (Figure 1I). Note that somewhat decreased 465 nm absorbance was observed upon completion of the reaction, which is likely attributable to pH-dependent variations in product absorptivity. Similar to as described above, complete consumption of DAB was confirmed via the addition of fresh catalyst and H_2_O_2_, which like before, did not result in any further increase in absorption (data not shown).

The DAB oxidation efficiency of Fe-TAML was also compared directly to that of HRP. In these analyses, we measure DAB oxidation using solutions containing different amounts of H_2_O_2_ (either 1 mM or 40 mM H_2_O_2_). The rationale for evaluating these conditions was based on our expectation that Fe-TAML would be less prone to inactivation by elevated levels of H_2_O_2_[16]. DAB oxidation by HRP was accelerated by a factor of 86-fold compared to that which was catalyzed by Fe-TAML, when carried out in pH 8.6 bicine buffer (Figure 1H). Nonetheless, due to Fe-TAML’s versatility as an oxidation catalyst, combined with advantages due to its small size, we proceeded to test Fe-TAML in cell-based DAB precipitation for EM imaging. Under these conditions, similar DAB yields could be achieved using hour long reaction conditions, as opposed to minutes, or by using multiple catalysts (note that 10 Fe-TAMLs could comprise only 10% of the mass of a single HRP complex).

### Cellular DNA labeling with Fe-TAML catalyzed DAB oxidation

Having confirmed the ability of Fe-TAML to catalyze DAB oxidation, we next investigated its utility in cytochemical labeling for EM. Here, we applied a biorthogonal ligation chemistry approach analogous to our previously reported “Click EM” strategy[17]. To do so, a click-reactive derivative of Fe-TAML was synthesized and the probe was reacted with cells labeled with metabolic DNA labeling substrates. An azide-modified Fe-TAML was synthesized from a previously reported alkyne [14] by a one-pot copper(I)-catalyzed azide-alkyne cycloaddition (CuAAC) reaction with 3-azido-(PEG)_2_-amine followed by reaction of the introduced amino group with 2-azidoacetate succinimidyl ester (Figure S1). Ion-pairing HPLC purification at pH 7 was necessary to avoid the loss of Fe^3+^ from the product, Fe-TAML azide (Figure 1D) that occurred with conventional acidic elution solvents.

Replicating HEK293T cells were labeled via an overnight pulse with EdU [18] and labeled DNAs were conjugated to Fe-TAML azide. The CuAAC reaction was carried out by subjecting fixed, permeabilized cells to a reaction mixture containing the probe and CuSO_4_. Freshly prepared ascorbate was added to the mixture to generate the active Cu(I) species *in situ* and the cells were reacted for 60 min with additional ascorbate added after 30 min. Following the reaction, excess reaction components were washed from cells prior to incubation with DAB and H_2_O_2_ for polymer deposition in bicine buffer pH 8.3, and deposition of optically dense polymer in the nucleii was visible by transmitted light microscopy within 2-3 minutes of incubation. Note that this reaction time is comparable to previous light-mediated DAB oxidation using EdU labeled cells conjugated with the photo-oxidizing dye DBF-azide[17]. Thus, although our solution-based analyses indicated an attenuated reaction rate for Fe-TAML compared to HRP, such rate did not require substantial procedural alterations to standard protocols. The generated DAB precipitates were confined to the nuclear compartment and enhancement with OsO_4_ staining further confirmed the nuclear specificity of DAB polymer formation. Note that signals from DAB precipitates colocalized with intensities from a conjugated AF647-azide, which was included within the CuAAC mixture as a fluorescence marker of EdU-labeled DNA (Figure 2A, B).

**Figure 2.**
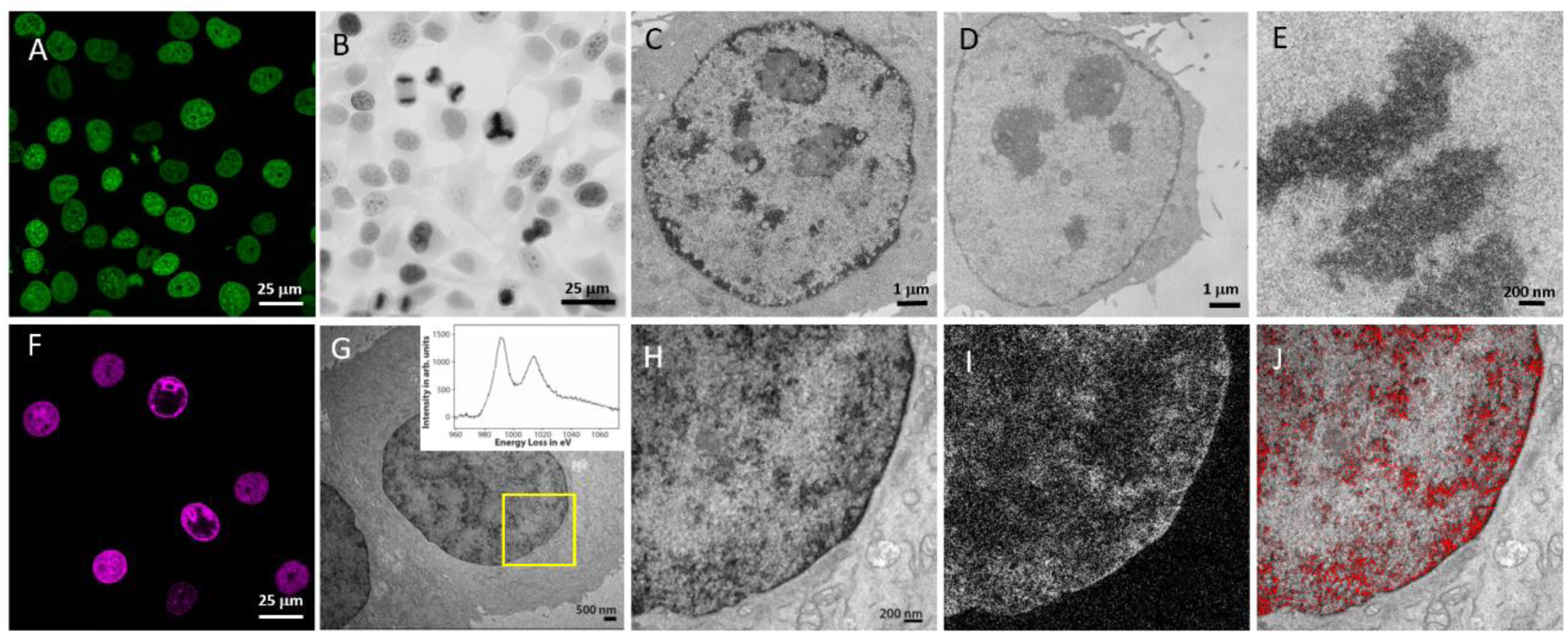
DNA labeling by click-EM with Fe-TAML azide. A) Fluorescent image of fixed HEK293 cells incubated with EdU followed by click reaction with AF488-azide. B) Transmitted light image of fixed EdU-treated cells clicked with Fe-TAML azide and then incubated with H_2_O_2_ followed by OsO_4_. C) TEM image of labeled cell nucleus. D) Control cell without reaction with Fe-TAML azide. E) High mag TEM of condensed chromatin. F-J) EELS and EFTEM to confirm specific labeling of Neodymium from Fe-TAML in SEA cells. F) Fluorescent image of combined Cy5-azide/Fe-TAML azide labeling. G) TEM image of cell after oxidation of Nd-DAB2 DNA obtained at a magnification of 6k (pixel size of 1.5 nm/pixel) with inserted EELS spectrum showing the presence of a strong (Nd) signal at region shown by box in TEM. The intensity unit is arbitrary. H) Expansion of boxed region in G. I) Nd elemental map obtained by EFTEM at a magnification of 6K and hardware binning of 4 pixels. J) Overlay of the Nd elemental map (red pseudocolored) on the conventional image.

After embedding in resin and sectioning to ∼100 nm, transmission EM (TEM) revealed patterns consistent with the staining of chromatin and the localization of newly synthesized DNA. High magnification inspection of cells captured during S- or G2 cell cycle phases revealed localized intensities at the nucleolar boundaries as well as circumferential staining along the nuclear lamina (Figure 2C). The specificity of the observed signals was especially apparent upon comparison with nearby cells that lacked nuclear contrast (Figure 2D), presumably due to the asynchronous nature of the culture, in which only a fraction of cells were undergoing active replication during the pulse with EdU. This result verifies the labeling specificity of the approach, demonstrating both the selectivity of Fe-TAML azide conjugation as well as its ability to produce high quality and tightly localized EM contrast surrounding its intended molecular target (in this case, newly replicated DNA). Dense mitotic chromosomes are clearly visible in cells undergoing prometaphase and metaphase (Figure 2E). The staining density and subcellular location of Fe-TAML catalyzed DAB oxidation is comparable to that achieved with click-EM[17]. In this method photosensitizing fluorophores containing azido groups were clicked to EdU-labeled cells and DAB is oxidized by intense illumination in the presence of O_2_.

### Multicolor EM with Fe-TAML

We then tested whether Fe-TAML can catalyze the oxidation and deposition by H_2_O_2_ of Ln-DAB2 which can be uniquely distinguished via elemental mapping in “multicolor-EM”. In this method[19], lanthanide ions bound to a precipitated DAB-DTPA chelate are selectively identified in an element specific manner by electron-energy loss spectroscopy (EELS) and with visual 2D mapping that can superimposed upon conventional contrast images via energy-filtered TEM (EF-TEM). Sequential deposition of distinct DAB-bound lanthanides can be used to generate 2-color elemental maps in which the localizations of separate orthogonally tagged species can be simultaneously overlaid on the conventional TEM. As a testbed to evaluate the suitability of Fe-TAML in EELS based multicolor EM, we conjugated the probe to EdU-fed telomerase-expressing primary small airway epithelial cells (hTERT-SAEC). Upon H_2_O_2_ mediated deposition of a Nd-DAB2 chelate we observed the expected nuclear optical densities, consistent with that which was observed above using conventional DAB and EdU-stained HEK293T cells as described above (Figure 2F, G). EM imaging of thin sections with spatially mapped EELS analysis confirmed the co-deposition of Nd at sites containing electron-dense (OsO_4_-stained) DAB polymers. Note that the specificity of Nd deposition was confirmed via detection of elemental signatures[20], evident through peaks at 990 and 1015 eVs (Figure 2G). The Nd elemental map (Figure 2H) pseudocolored overlay with the TEM is shown in Figure 2I and high intensities are associated with corresponding dense osmium staining in the TEM image, in contrast to visible cytoplasmic organelles such as mitochondria.

Having verified the compatibility of Fe-TAML and Nd-DAB2, we next examined its utility in orthogonal multicolor EM labeling of distinctly labeled targets. Here, we used hTERT-SAEC transduced with adenovirus encoding the viral E4-ORF3 protein (which forms nuclear fibrils) fused with the genetically encoded photooxidizing EM probe miniSOG (E4-ORF3-miniSOG) [21,22]. To introduce a clickable label to which Fe-TAML azide could be attached we labeled these cells with EdU and carried out CuAAC reactions as described above. Sequential elemental labeling was then carried out using distinct lanthanide-DAB reagents. First, Ce-DAB2 was deposited via photoexcitation of miniSOG using illumination with blue light (450-490 nm). Labeled nuclear fibrils characteristic of E4-ORF3 polymerization are visible in the transmitted light image (Figure 3B) that closely correlate with the image of miniSOG fluorescence collected before photooxidation (Figure 3A). After removal of unreacted reagents, reactive amines on the deposited Ce-DAB were blocked using acetylimidazole, to prevent their further polymerization with distinct DAB species during further deposition steps. Cells were then subjected to CuAAC ligation with Fe-TAML azide (in combination with dilute Cy5-azide as a fluorescence co-marker) and subsequent H_2_O_2_ mediated oxidation was carried out using Nd-DAB2 in a bicine buffered solution of pH 8.3. (Figure 3D). Additional nuclear staining is now visible in the transmitted light image that co-registers with the bright fluorescent Cy5 staining (Figure 3C). EELS spectra and an overlay of the Nd and Ce elemental maps with the TEM of such a doubly stained region confirmed the distinct spatial distributions of the viral protein and cellular DNA (Fig. 3E-H). Worthy of note, Ce-DAB2 (marking E4-ORF3-miniSOG) and Nd-DAB2 (marking EdU-labeled DNA) were colocalized in an intertwined-manner within discrete sub locales of the nuclear matrix, an observation that agrees with the previously described heterochromatin assemblies that surround E4-ORF3 polymer chains [21]. Together, these results validate the utility of Fe-TAML as a sensitive and orthogonal multicolor EM marker that can be combined with miniSOG to visualize distinct biomolecular species.

**Figure 3.**
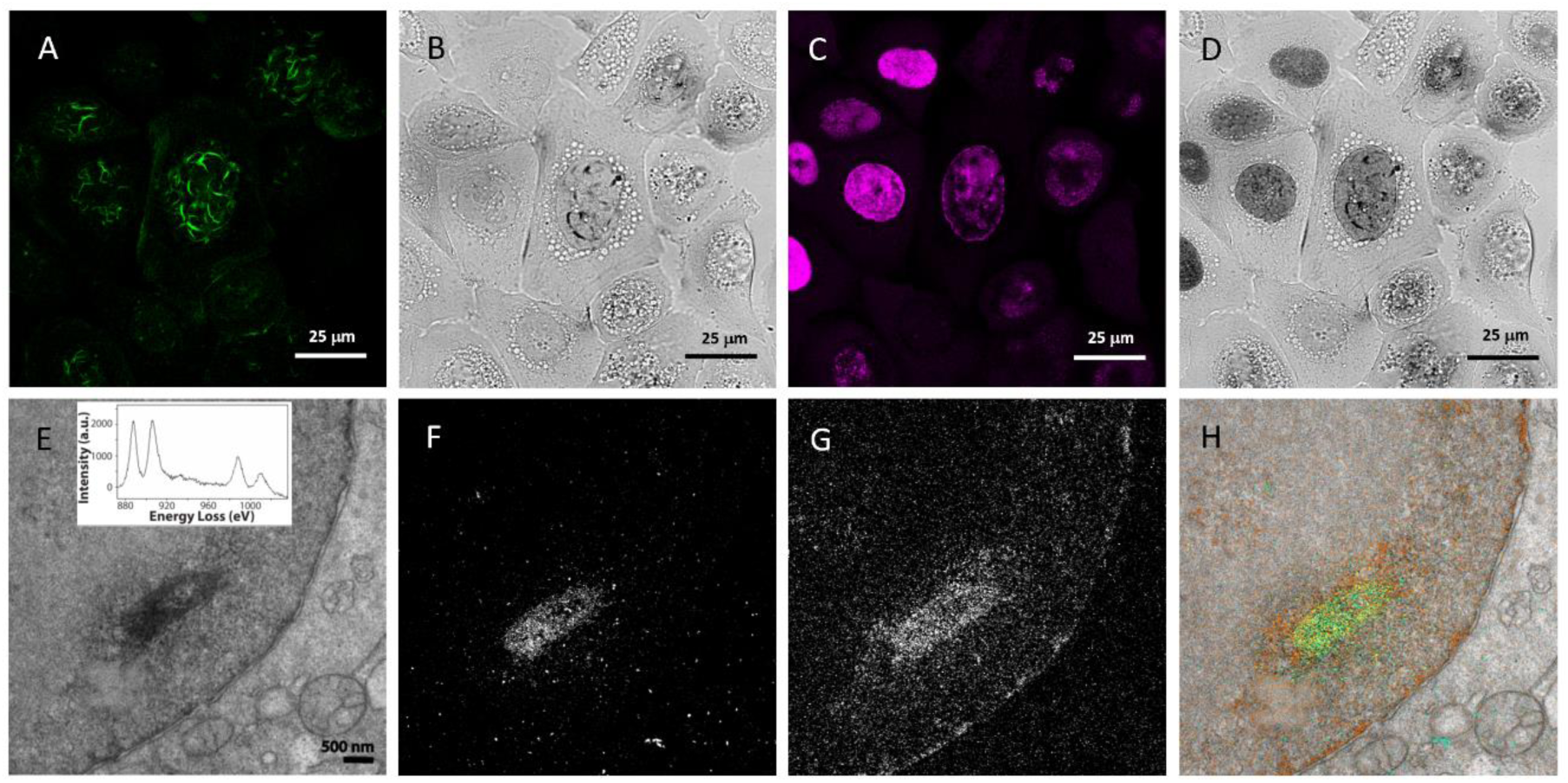
Two color EM of EdU-treated SEA cells transduced with E4-ORF3-miniSOG and DNA labeled with Fe-TAML azide and Cy5-azide. A) Fluorescent confocal image of miniSOG labeling. B) Transmitted light image of the same field of cells in (A), after miniSOG photooxidation with Ce-DAB2. C) Fluorescent image of subsequent Cy5 azide and Fe-TAML azide labeling of DNA in same field of cells. D) Transmitted light image of cells after Fe-TAML oxidation of Nd-DAB2 with H_2_O_2_. E) Convention TEM image of cell nucleus and cytoplasm from cells treated as in (D) with inset of background subtracted EELS spectrum showing Ce and Nd core-loss edges. F) Correlated Ce elemental map obtained by EFTEM. G) Correlated Nd elemental map obtained by EFTEM. H) Pseudocolor overlay of Ce (green) and Nd (red) elemental maps over conventional TEM image (E).

### Genetically targeted Fe-TAML

In addition to examining metabolically labeled structures we also investigated the utility of Fe-TAML in visualizing individual protein species. In this approach, we applied self-labeling protein tags including HaloTag[5] and SNAP-tag[4]. These are widely used tags that can react to form covalent conjugates with synthetic chloroalkane (CA) or benzylguanine (BG) substituent respectively. Genetically encoded fusions with these tags can be labeled and imaged in cells or *in vivo* by incubation with membrane permeant fluorescent ligands. To test the suitability of protein-specific labeling, we generated a trifunctional probe, CA-Fe-TAML-rhodamine containing Fe-TAML functionalized with a chloroalkane handle (for HaloTag targeting) and linked with a rhodamine dye (for direct fluorescence visualization). This probe was synthesized from Nα-Fmoc-S-trityl-L-cysteine by reaction with 3-azidopropylamine to form the amide at the free carboxyl group, followed by Fmoc deprotection and reaction of the N_α_-amine with 5(6)-carboxytetramethylrhodamine succinimidyl ester (Figure S2). The S-trityl ether was subsequently cleaved with TFA and resulting free thiol reacted with a 2-pyridyl-dithiopropanoyl amide of HaloTag linker amine. Finally, CuAAC reaction of the azido group with Fe-TAML alkyne gave the desired product, CA-Fe-TAML-rhodamine (Figure 1C) that was purified by reverse phase HPLC at pH 7 and characterized by mass spectroscopy. Covalent attachment to purified HaloTag protein following incubation was confirmed by LC-MS with the expected increase in mass of 1357.9 indicating loss of Cl by reaction with active site Asp106 and loss of Fe following HPLC under acidic conditions. The time course for reaction of CA-Fe-TAML-rhodamine was rapid occurring within 1 minute after mixing and giving over 2-fold increase in fluorescence upon binding (Figure S4A). Similar fluorogenic behavior of related fluorophores on reaction with HaloTag protein has been widely reported[23]. The fluorescence quantum yield for bound CA-Fe-TAML-rhodamine was determined to be 0.12 with excitation and emission maxima at 555 nm and 582 nm respectively (Figure S4B).

Labeling permeabilized and fixed human osteosarcoma (HOS) cells stably expressing histone H2B-HaloTag with CA-Fe-TAML-rhodamine gave brightly stained fluorescent nuclei (Figure 4A). Note that initial live cell treatments with the probe did not produce such fluorescent patterns, which suggested the requirement of slight membrane permeabilization for efficient intracellular probe delivery. Importantly, control (non-transfected) cells were non-fluorescent upon permeabilization and staining with the probe (Figure 4C), thus confirming its labeling specificity against HaloTagged proteins. DAB oxidation by H_2_O_2_ at pH 8.3 of stained cells gave nuclear osmiophilic precipitates consistent with the expected subcellular localization of histone proteins across the cell cycle including interphase, telophase and clearly defined mitotic chromosomes in anaphase (Figure 4B) compared with untransfected HEK23T cells that show minimal fluorescence. Subsequent higher magnification with TEM confirmed the nuclear distribution and morphology of the precipitated DAB (Figure 4D), and high-resolution 3D tomographic analyses revealed the resolution quality of the probe design, with which individual nucleosomal puncta were visible with the expected molecular dimension, including discs of 11 nm in diameter and of 5.5 nm depths (Figure 4E-H). A 3-dimensional rendering of a nuclear region in the 100 nm slice is shown in Figure 4H. Note that the labeling specificity and resolutions that was achieved via CA-Fe-TAML-rhodamine is comparable with those which have been reported using JF570-CA[24]. Although labeling procedures with JF570 report somewhat shorter reaction times (7 min) as compared with our tests with Fe-TAML, an advantage of our approach is H_2_O_2_ mediated DAB labeling can be performed via timed bench-side reactions, with monitoring by standard transmitted light microscopes and without the need for establishing on-scope staging for photo-oxidation. In addition, we anticipate that a SNAP-tag JF570 reactive probe could be combined with CA-Fe-TAML-rhodamine for dual lanthanide labeling via EELS as described above. Overall, our results validate the utility of CA-Fe-TAML-rhodamine in protein-specific CLEM labeling.

**Figure 4.**
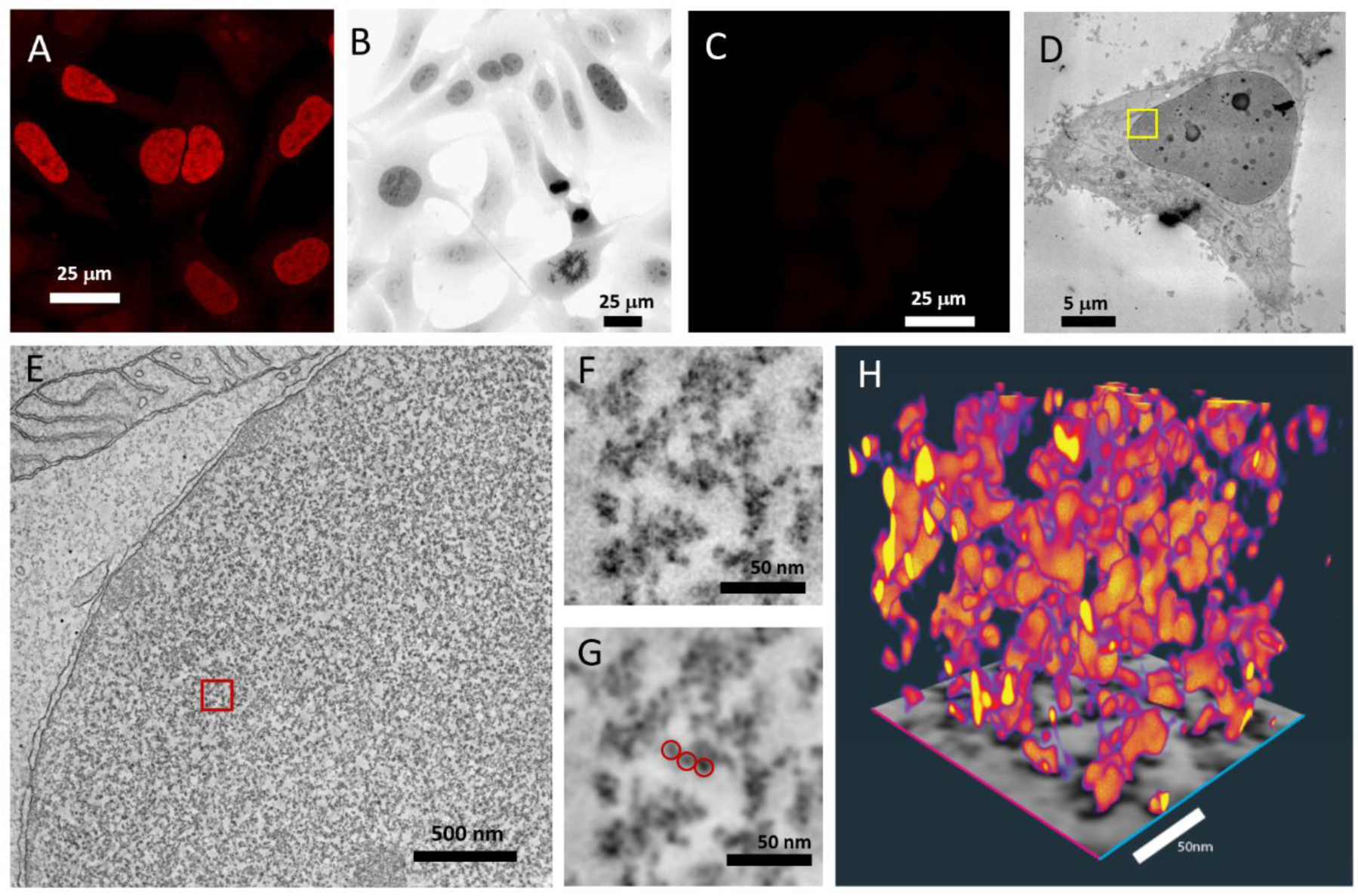
Imaging of HOS cells stably or HEK293T cells transiently expressing histone H2B-Halotag followed by labeling with CA-Fe-TAML-rhodamine and oxidation of DAB. A) Confocal fluorescence image of HOS whole nuclei stably expressing H2B-Halotag after labeling with CA-Fe-TAML-rhodamine. B) Bright-field image of labeled HOS cells as in (A), after DAB oxidation and osmium staining. The field of view show cells at different mitotic stages. C) Confocal fluorescence image of CA-Fe-TAML-rhodamine labeled non-transfected HEK293 cells. D) TEM micrograph of a HEK293 cell expressing HaloTag–H2B fusion proteins labeled with CA-Fe-TAML-rhodamine, followed by oxidation of DAB and osmification. E). Electron tomogram slice of the region of interest highlighted by the yellow box shown in (D). F) Enlarged view of the area delimited by the red box in (E). G) Visualization of the same area of the tomogram slice shown in (F) after anisotropic filtering. Circles (11 nm diameter) highlight a cluster of nucleosomes in the volume. H) 3D volumetric rendering extracted from the electron tomogram in (E-G).

### Immunocytochemical labeling of endogenous proteins with Fe-TAML

Lastly, we tested the utility of Fe-TAML as a probe for detecting endogenous (untagged) protein targets via immunolabeling with Fe-TAML conjugated antibodies. Indeed, immunolabeling methods remain a mainstay in the histochemical, fluorescence, and EM detection of proteins in diverse specimens. Recognizing the utility of Fe-TAML in immunolabeling, where the decreased size of Fe-TAML could prove advantageous, we generated Fe-TAML-conjugated secondary antibody and utilized the probe to visualize assembled microtubules in cells stained with a primary anti-beta-tubulin IgG. In order to generate the Fe-TAML-labeled secondary we reacted Fe-TAML alkyne to goat-anti mouse immunoglobulin that had been labeled on multiple reactive lysine residues or reduced interchain cysteine disulfides with an azido bifunctional crosslinkers containing either a N-hydroxysuccinimide (NHS) ester for lysine modification or a maleimide for cysteines. To enable correlative imaging with these antibodies they were also conjugated with Cy5-NHS. The subsequent CuAAC reaction with Fe-TAML alkyne followed by gel filtration purification gave Fe-TAML/Cy5 secondary IgG that was then tested on fixed HeLa cells that had been incubated with a monoclonal primary antibody to β-tubulin. We varied the number of Fe-TAML attached to the IgG and examined both binding, fluorescence and DAB catalyzed oxidation with optimal results obtained with approximately 10 Fe-TAML and one Cy5 label per antibody. Labeling stoichiometries were determined by absorbance of the Fe-TAML and Cy5 moieties at 365 and 650 nm respectively. Reaction via lysine rather than cysteine residues was preferable as no prereduction step was required, azido-peg-NHS crosslinkers are commercially available and stable unlike the corresponding azido-peg-maleimide versions and more than 8 Fe-TAML could be conjugated as IgG contains only 4 solvent exposed disulfide bonds.

A fluorescent image of immunolabeled β-tubulin with the Fe-TAML/Cy5 conjugated secondary antibody in HEK293 cells is shown in Figure 5A. Clearly defined microtubule bundles are visible in quiescent cells and during stages of the cell cycle including the mitotic spindle and telophase. Individual microtubules visible in less dense areas demonstrate the high specificity of labeling with the Fe-TAML/Cy5 conjugated secondary antibody. DAB oxidation was carried out at pH 8.3 for 60-90 min at 4°C, and after post-fixation and osmification, distinct labeling of the microtubules is visible by bright field light microscopy (Figure 5B). Following embedding and sectioning, TEM revealed that the microtubules are distinctly though lightly stained with electron-density and visibly distinguishable from cytoplasmic material (Figure 5C). Higher magnification images of microtubule bundles and single microtubules are shown in Figure 5D and E respectively. The fluorescent staining with Fe-TAML/Cy5 is similar to that obtained with a commercial FITC-conjugated secondary antibody (Figure 5F) and only weak non-specific labeling or autofluorescence is seen when no primary antibody is present (Figure 5G, H). Incubation of these controls with DAB and H_2_O_2_ gave no fibrillar staining (Figure 5I, J). The retention of antibody binding despite the higher labeling levels of Fe-TAML compared to those generally used for fluorochromes may reflect its charged, more hydrophilic structure and inclusion of a short peg linker.

**Figure 5.**
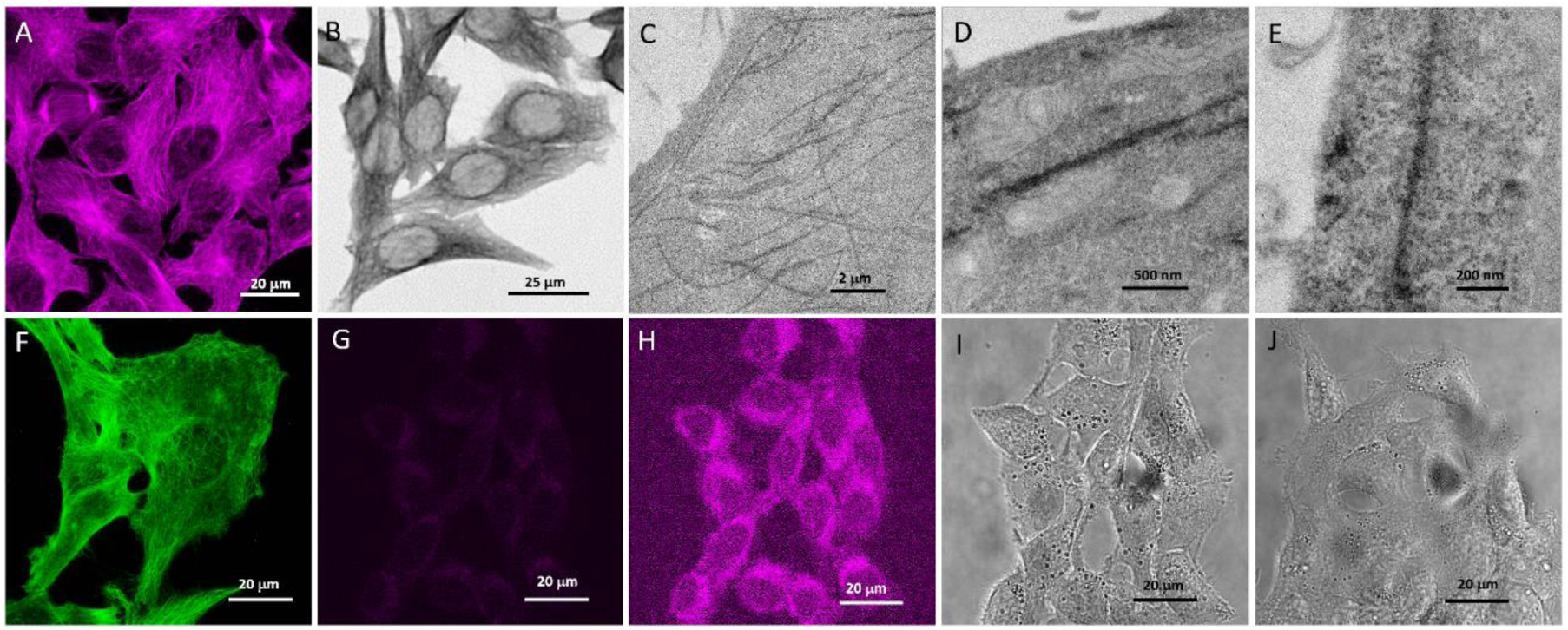
Immunolabeling of microtubules with Fe-TAML/Cy5 conjugated antibody in HEK293 cells. A) Confocal fluorescence image of cells labeled with mouse anti-β-tubulin monoclonal antibody and secondary Fe-TAML/Cy5 conjugated goat anti-mouse secondary. B) Bright-field image of osmium-stained cells in A after oxidation of DAB by H_2_O_2_ catalyzed by Fe-TAML. C) Electron micrograph of cell from B with specific dark staining of microtubules. D) Higher magnification TEM image of microtubule bundle. Note the adjacent mitochondria with well-preserved cristae. E) Higher magnification TEM image of single stained microtubule. F) Confocal fluorescence image of cells labeled as in A but with FITC-conjugated secondary antibody. G) Confocal fluorescent image of cells labeled in A but with no primary anti-tubulin antibody using same instrument settings. H) Brightened image of G to illustrate dim background staining and autofluorescence without primary antibody. I) and J) cells from F) and G) respectively after incubation with DAB and H_2_O_2_ and resulting bright-field image with no visible microtubule staining.

The diameter of Fe-TAML oxidized DAB precipitate coating individual MTs is ∼50-75 nm consistent with their known diameter of ∼25 nm plus two attached antibodies (each IgG is 12 nm). Fe-TAML stained MTs are comparable to that generated by secondary antibodies labeled with the photosensitizer, eosin[25]. Photosensitized oxidation of DAB requires brief (few minutes) of intense localized irradiation in the presence of saturating oxygen while monitoring the progression of precipitation by intermittent brightfield microscopy. By comparison, staining with APEX2 with a N-terminal fusion to α-tubulin restricts oxidation to the enclosed inner core of the MT[26]. Restriction of DAB oxidation products and intermediates is aided by the dense mesh produced by post-labeling fixation with 2% glutaraldehyde that also optimizes preservation of cellular ultrastructure in the TEM. Overall, these results confirm the utility of TAML-conjugated antibodies as versatile and high-resolution labeling probes for visualizing endogenous (untagged) cellular structures.

## SUMMARY

In this work, we investigate the use of the small molecule peroxidase, Fe-TAML for correlated light and electron microscopy and color EM[19], demonstrating its successful application for marking DNA by click-EM[17], expressed proteins with a HaloTag label, and for immunolabeling of endogenous proteins. Compared to the protein peroxidases, HRP and APEX2, Fe-TAML has significantly slower catalytic rates for DAB oxidation at physiological pH that can be partially overcome by working at more alkaline conditions following fixation and with higher concentrations of H_2_O_2_. The slower rates can be an advantage allowing finer control of reaction progress whereas using HRP and Apex2 often requires precise timing (measured in seconds) with constant monitoring of the optimal faint darkening followed by a rapid washout of H_2_O_2_, to prevent over reaction. The Fe-TAML peroxidase reaction appears to give comparable sensitivity and resolution to our previous efforts for marking DNA to DAB photooxidation by dibromofluorescein using click-EM[17], and by the intercalator dye, DR in ChromEM[27]. We also see a similar quality of labeling for EM of H2B histone HaloTag with Fe-TAML to that we demonstrated with JF570 photosensitizer[24] with a high sensitivity of detection of the 1-2 molecules of H2B-Halo in each labeled nucleosome. Immunolabeling with Fe-TAML gave specific and highly spatially restricted coating of microtubules similar to our previous work using eosin-mediated photooxidation of DAB[28].

The small size and hydrophilicity of Fe-TAML permits labeling antibodies for immunohistochemistry with multiple (∼10) Fe-TAML with a minimal increase in size while maintaining antigen binding. This contrasts with the fluorophores used in immunofluorescence where self-quenching and hydrophobicity limit such high labeling stoichiometries. This will also facilitate greater penetration into fixed tissue than conventional IHC using biotinylated antibody with Streptavidin-HRP (∼100kD), nanometer-sized immunogold, or Quantum Dots that require milder fixation conditions resulting in poor cellular ultrastructure with EM.

Fe-TAML, like APEX2 and HRP is not fluorescent but a wide range of fluorophores covering the entire spectrum can be readily incorporated with a minimal increase in molecular weight (total <1.5kD) for verification of labeling and CLEM. Fusions of fluorescent proteins with APEX and HRP[8] are substantially larger (about 54kD and 71kD respectively). As we demonstrate, for labeling DNA/RNA by click-EM and for immunofluorescence, co-labeling with a small molecule fluorophore is possible, again minimizing probe size. In our designs we chose Cy5 and TMR as dual labels with Fe-TAML. The rationale for these selections was on the basis of limiting any light dependent DAB oxidation. Thus, in future applications, especially for 2 color labeling, users should consider one of these dyes (or another appropriate non-photo-oxidizing fluorophore) when implementing. Fe-TAML is currently limited for use with fixed cells, but increased membrane permeability should be achievable with some chemical modification enabling live cell labeling of HaloTag or SNAP-tag fusions. Additional chemical modifications of Fe-TAML to increase catalytic rates have been reported[12] although with tradeoffs in stability and therefore turnover number[29]. Proximity labeling with conjugates of Fe-TAML may be possible and applicable to small biomolecules and ligands unlike available protein-based fusions[30].

## MATERIALS AND METHODS

### Materials

Reagents were from Sigma-Aldrich (St. Louis, MO), solvents and cell culture reagents were obtained from Thermo Fisher Scientific (Pittsburgh, PA) except where noted. Fe-TAML B* and Fe-TAML alkyne triethylammonium salts were obtained from (GreenOx Catalysts Inc., Pittsburgh, PA) as custom syntheses.

Reactions were monitored by LC-MS (Ion Trap XCT with 1100 LC, Agilent, Santa Clara, CA) using an analytical Luna C18(2) reverse-phase column (Phenomenex, Torrance, CA), acetonitrile/H2O (with 0.05% v/v CF_3_CO_2_H) linear gradients, 1 mL/min flow, and ESI positive or negative ion mode. PLRP-S columns (Agilent) were used for protein conjugates with similar acetonitrile/water/0.05% TFA gradients. For ion pairing HPLC, linear gradients of acetonitrile/0.1M triethylammonium acetate (TEAA) buffer pH 7 were used on analytical Luna C18(2) columns. Compounds were purified by preparative HPLC using the above gradients and semi-preparative Luna C18(2) columns at 15 ml and 3.5 ml/min respectively. UV−vis absorption spectra were recorded on a Shimadzu UV-2700 (Kyoto, Japan) spectrophotometer and fluorescence spectra on Fluorolog 3 (Horiba Scientific, Piscataway, NJ).

DNA constructs.

For the construct of HaloTag-H2B (JH1348), the H2B sequence was amplified from pminiSOG-H2B-6[31] and substituted EGFP in pHaloTag-EGFP (addgene #86629)[32] through the In-Fusion cloning method.

## EXPERIMENTAL PROCEDURES

### Fe-TAML azide

To a solution of Fe-TAML alkyne triethylammonium salt in DMSO (12.5 μmol, 50 mM, 250 μL) in a plastic screw cap tube, the following solutions were added in the order listed; Na-MOPS pH 7.2 (100 μmol, 1 M solution in water, 100 μL), pre-mixed CuSO_4_ (1.25 μmol, 10 mM in water, 125 μL) and THPTA (2.5 μmol, 10 mM in water, 250 μL), 11-azido-3,6,9-trioxaundecan-1-amine (90%, 13.5 μmol, 100 mM in DMSO, 150 μL; Fluka) and placed under an Ar atmosphere. Freshly prepared sodium ascorbate (25 μmol, 100 mM in water, 250 μL) was added and the reaction mixture was mixed and kept at room temperature for 1 hr when LC-MS revealed complete reaction. Azidoacetic NHS ester (37.5 μmol, 100 mM in DMSO, 375 μL, Click Chemistry Tools) was added with mixing and left overnight at room temperature when LC-MS revealed reaction is complete. The desired product was separated by RP HPLC using a TEAA pH 7-ACN 5-90 % gradient in 20 min, lyophilized to an orange oil, and dissolved in DMSO (200 μL) to give a 14 mM solution by absorbance at 365 nm using an extinction coefficient of 5800 M^-1^cm^-1^. Yield 22%, 86% pure (side product mass 1183) by RP-HPLC; ES-MS (m/z) [M]^+^ for C_34_H_50_N_12_O_9_ 771.4 following loss of Fe^3+^ under acidic conditions; found 771.4. ES-MS (m/z) [M]^-^, MeOH) for C_34_H_46_FeN_12_O_9_, 822.3; found 822.2.

### CA-Fe-TAML-rhodamine

#### NH_2_-cys(S-Trt)-CO-NH-(CH_2_)_3_-N_3_

Fmoc-NH-cys(S-Trt)-OH (19.8 mg, 33.8 μmol) and HATU (14.1 mg, 37.2 μmol) were dissolved in dry DMF (100 µL) in a plastic screw cap tube and 3-azidopropyl-1-amine (3.7 μL, 37.2 μmol, Click Chemistry Tools) followed by DIEA (13 μL, 74.42 μmol) were added with mixing. The reaction mix turned yellow and LC-MS revealed complete reaction in 30 mins; ES-MS (m/z) [M+ Na]^+^ for C_40_H_37_N_5_NaO_3_S, 690.25; found 689.3. Piperidine (20 μL, 0.2 mmol) was added and the solution evaporated under high vacuum after 1 h, dissolved in DMSO, separated by RP-HPLC and lyophilized to give a white solid. Yield, 14.3 mg, 76%. ES-MS (m/z) [M]^+^, [M+ Na]^+^ for C_25_H_27_N_5_OS, 446.2, 468.2; found 446.1, 468.1.

#### 5(6)-TMR-CONH-cys(SH)-CO-NH-(CH2)3-N3

5(6)-Carboxytetramethylrhodamine, 5(6)-TMR-CO_2_H (1.35 mg, 3.14 μmol, Novabiochem) and TSTU (1.3 mg, 4.4 μmol) were dissolved in dry DMSO (25 μL) with TEA (0.96 μL, 6.9 μmol) and kept at room temperature. Reaction was complete in 30 min (by LC-MS) and then added to a solution of NH_2_-cys(S-Trt)-CO-NH-(CH_2_)_3_-N_3_ (2.0 mg, 3.6 μmol) in DMSO (10 μL) with NMM (1 μL, 9.1 μmol) and kept at room temperature overnight when LC-MS revealed complete reaction. After acidification with HOAc (2 μL), the desired product was isolated by RP-HPLC and lyophilized to a red solid. Yield, 2.0 mg (74%) ES-MS (m/z) [M]^+^ for C_50_H_48_N_7_O_5_S, 858.3; found 858.3. The trityl group was removed by dissolving the product (1.89 mg, 2.2 μmol) in TFA:H_2_O:Triisopropylsilane:Ethanedithiol (92.5:2.5:2.5:2.5 v/v, 0.5 mL) for 30 mins, evaporation under high vacuum, purification by RP-HPLC and lyophilization to a red solid. Yield, 0.9 mg (66%) ES-MS (m/z) [M]^+^ for C_31_H_34_N_7_O_5_S, 616.2; found 616.1.

#### SPDP-HaloTag linker

HaloTag linker amine[33] (3.0 mg, 13.5 μmol) and SPDP (4.7 mg, 15 μmol) were dissolved in dry DMSO (50 μL) and NMM (3.3 μL, 30 μmol) added. LC-MS revealed reaction was complete after overnight when the reaction mixture was neutralized with HOAc (5 μL) and the product was purified by RP-HLPC (and lyophilized to a colorless oil. Yield, 3 mg (54%) ES-MS (m/z) [M]^+^ for C_18_H_29_ClN_2_O_3_S_2_, 420.1; found 421.1.

#### 5(6)-TMR-CONH-cys(S-S-propanoyl-HaloTag ligand)-CO-NH-(CH_2_)_3_-N_3_

A solution of 5(6)-TMR-CONH-cys(SH)-CO-NH-(CH_2_)_3_-N_3_ in DMSO (100 μL, 6.25 mM measured by absorbance in 0.1 M HCL in 95% ethanol using εmax 95000 M^-1^cm^-1^ at 554 nm, 0.626 μmol) was mixed with SPDP-HaloTag linker (20 μL, 49 mM in dry DMSO, 0.98 μmol) and NMM (1 μl, 10 μmol) added. After 1 h, HOAc (5 μL) was added and the desired product purified by RP-HPLC to give a colorless oil. Yield, 0.42 mg (72%) ES-MS (m/z) [M]^+^ for C_44_H_58_ClN_8_O_8_S_2_, 925.4; found 925.4.

#### CA-Fe-TAML-rhodamine

To a solution of 5(6)-TMR-CONH-cys(S-S-propionyl-HaloTag ligand)-CO-NH-(CH_2_)_3_-N_3_ (50 μL, 4.5 mM in DMSO; 0.225 μmol) was added sequentially Na-MOPS (20 μL of 1 M in water, pH 7.2), Fe-TAML alkyne triethylammonium salt (60 μL, 5 mM solution in DMSO, 0.30 μmol), premixed CuSO_4_ (100 μL, 10 mM in water, 1 μmol) and THPTA (100 μl, 50 mM in water, 5 μmol). The solution was placed under Ar and freshly prepared sodium ascorbate (100 μL, 100 mM in water, 10 μmol) was added, the reaction mixture was mixed and kept at room temperature overnight when LC-MS (ACN-H_2_O-TFA) revealed complete reaction (ES-MS (m/z) [M]^+^ for C_68_H_89_ClN_13_O_13_S_2_, 1394.6 (-Fe^3+^); found 1394.4. The desired product was separated by RP-HPLC using a TEAA pH 7 - ACN gradient, lyophilized to an orange oil, and dissolved in DMSO (200 μL) to give a 0.4 mM solution by absorbance at 554 nm using an extinction coefficient of 92000 M^-1^cm^-1^ in 0.1 M HCl-95% ethanol. Yield, 35% ES-MS (m/z) [M]^-^ for C_68_H_85_ClFeN_13_O_13_S_2_, 1445.5; found 1445.2.

#### Reaction with HaloTag protein

CA-Fe-TAML-rhodamine (15 μM) was reacted with 30 μM his_6_-tagged HaloTag protein (ES-MS (m/z) [M]^+^; found 34795.8 deconvolved) in 0.1M Na MOPS pH 7.2 for 1 h and the adduct was detected by RP-HPLC (Agilent PLRP-S column), ES-MS (m/z) [M]^+^; found 36153.7. Mass difference for addition of C_68_H_87_N_13_O_13_S_2_ (-HCl, -Fe), 1357.6; found 1357.9.

The fluorescence quantum yield of bound CA-Fe-TAML-rhodamine to HaloTag protein was 0.12 as determined in the above complete reaction by comparison with rhodamine B in EtOH (QY=0.68)[34]

### Fe-TAML-peg4-Cy5 goat anti-mouse IgG conjugate

Goat anti-mouse (GAM) IgG (100 μL 2.3 mg/ml, 1.53 nmol; H+L, Affinipure 115-005-003, Jackson ImmunoResearch) and 1 M-bicine buffer pH 8.3 (10 μL) were mixed with azido-PEG4-NHS (2.3 μL of a 10 mM freshly prepared solution in DMSO, 23 nmol, 15 eq; Click Chemistry Tools) at 4°C overnight. Cy5-NHS (1.2 μL of a 2.56 mM freshly prepared solution in DMSO, 3.06 nmol, 2 equiv; GE Healthcare) was added with mixing and kept at room temp for 4 h. The conjugate was purified by collecting the first blue colored band on gel filtration with Sephadex G-25 (0.6 g in PBS) and concentrated to 160 μL by ultrafiltration (30 kD MWCO Amicon Ultra, Millipore). UV-vis indicated 6.8 μM IgG and 5.3 μM Cy5 using extinction coefficients of 230000 and 12500 M^-1^cm^-1^ at 280 nm respectively and 250000 M^-1^cm^-1^ for Cy5 at 650 nm. 1M Na MOPS buffer pH 7.2 (16 μL) was added followed by a premixed CuSO_4_ (16 uL, 10 mM in water) and THPTA (16 μL, 50 mM in water) and then Fe-TAML alkyne (2.5 μL 5 mM in DMSO, 23 nmol, 15 eq.) Freshly prepared sodium ascorbate (8 μmol, 100 mM in water, 80 μL) was added and the reaction mixture was mixed and kept at room temperature overnight with protection from light. The product was separated by gel filtration (1 g Sephadex G-25 in PBS), collection of the first colored band and concentrated to 150 μL by ultrafiltration. UV-vis indicated 4.2 μM IgG, 42 μM Fe-TAML and 3.3 μM Cy5 using extinction coefficients of 5600 and 7500 M^-1^cm^-1^ at 365 nm for Fe-TAML and N3-peg4-Cy5 IgG respectively and assuming no loss of Cy5 in the click reaction. The conjugate was stored at 4°C and used within 7 days.

### Reaction kinetics of Fe-TAML B* and Fe-TAML alkyne catalyzed oxidation of diaminobenzidine by H_2_O_2_

50 mg gelatin (Kodak type) was added to water (25 mL) at 70°C with stirring until dissolved (5 mins) and cooled to room temp. Diaminobenzidine (5.4 mg free base, 25 µmol) was added with sonication to aid dissolution, centrifuged and the resulting decanted solution was diluted with an equal volume of either 0.2 M cacodylate pH 7.4, 20 mM bicine 0.2 M NaCl pH 8.3, or 20 mM bicine 0.2 M NaCl pH 8.6 to give final pH values of 7.41, 8.27 and 8.55 respectively. Reaction kinetics were determined in triplicate at room temperature (22°C) at 465 nm in cuvets by addition of Fe-TAML B*, Fe-TAML alkyne from stock solutions in dry DMSO with mixing. Freshly prepared H_2_O_2_ solutions were prepared from 30% H_2_O_2_. To determine Km values for DAB, appropriate volumes of a freshly prepared stock solutions in DMSO were added to the gelatin-containing buffer. Progress curves from initial rates (reaching absorbance values of 1) were fitted to linear fits (Prism, GraphPad Software, San Diego). Extinction coefficients for oxidized DAB at 465 nm were calculated from reactions allowed to proceed to completion when additional Fe-TAML and H_2_O_2_ caused no further absorbance increase.

Horse Radish Peroxidase (Pierce^TM^, 31490 Thermo-Fisher) concentration determined by the absorbance at 403 nm using an extinction coefficient of 109,000 M^-1^cm^-1^[35]

### Labeling of EdU-treated HEK293T cells with Alexa Fluor 488 azide

Alexa Fluor 488 azide labeling was accomplished as previously described[17]. HEK293T cells were plated onto MatTek dishes containing 35 mm glass bottoms No. 0 coverslips that were coated with poly-d-lysine. (P35GC-0-14C, MatTek Corporation). The cells were incubated with 10 μM EdU (1149-100, Click Chemistry Tools) for 12 hours. Cells were fixed with 2% glutaraldehyde (18426, Ted Pella Incorporated) in 0.1 M sodium cacodylate buffer, pH 7.4 (18851, Ted Pella Incorporated) containing 1 mM CaCl_2_**^.^**2H_2_O (223506-500G, Sigma-Aldrich) for 5 minutes at 37^°^C and then on ice for 55 minutes. Glutaraldehyde was removed from the cells, were washed with 0.1M sodium cacodylate buffer pH 7.4 (5 × 1 min) on ice, washed with PBS pH 7.4 without calcium and magnesium (21-040-CV, Corning) (2 × 1 min) at room temperature and rinsed 2 times with filtered (0.22 μm Millex 33mm PES sterile filter, SLGSR33RS, Sigma-Aldrich) freshly made 1% BSA (Sigma, A8022-100G) in PBS pH 7.4 (2 × 1 min) at room temperature. Click reaction of the cells was carried out at room temperature and protected from light by using a freshly prepared 1.0 μM Alexa Fluor 488 azide solution from 450 μl click buffer (50 mM Na·HEPES pH 7.6 (H-3375, Sigma), 100 mM NaCl (59625-5KG, Sigma-Aldrich), recipe), 5.0 μl CuSO_4_ (100 mM in water), 1 μl Alexa Fluor 488 azide (5.6 mM) and the reaction initiated with 50 *μ*l of freshly prepared aqueous sodium ascorbate (100 mM) (A7631-25G, Sigma-Aldrich) with a second 50 *μ*L aliquot added after 30 min for a total of 60 minutes. The cells were washed with filtered 1% BSA in PBS pH 7.4 (2 × 1min) at room temperature, and PBS pH 7.4 (5x 1 min) at room temperature. Fluorescence imaging of the labeled cells with Alexa Fluor 488 azide were acquired by a Leica SPEII confocal microscope.

### Labeling and DAB oxidation of EdU-treated HEK293T cells with Fe-TAML azide

HEK293T cells were plated onto MatTek dishes containing 35mm glass bottom No. 0 coverslips coated with poly-d-lysine. The cells were incubated with 10 μM EdU for 12 hours and fixed with 1% glutaraldehyde in PBS, 1X buffer, pH 7.4 for 5 minutes at 37^°^C and then on ice for 55 minutes. Fixative was removed and cells were washed with 1X PBS, 1X pH 7.4 (5 × 1 min) on ice. The cells were treated for 5 minutes with -20°C methanol (A452-1, Fisher). The methanol was removed, and cells were washed (3 × 1min) with cold PBS, pH 7.4 washed with PBS, pH 7.4 (2 × 1 min) at room temperature and rinsed (2 × 1 min) with filtered 1% BSA in PBS, 1X pH 7.4 at room temperature. Click reaction of the cells was carried out at room temperature and protected from light by using a 25 μM Fe-TAML azide solution freshly prepared from 450 μl click buffer (50 mM Na·HEPES pH 7.6, 100 mM NaCl), 5.0 μl CuSO_4_ (100 mM in water), 1 μl Fe-TAML azide (14 mM in dmso) and the reaction initiated with 50 *μ*l of freshly prepared aqueous sodium ascorbate (100 mM) with a second 50 *μ*L aliquot added after 30 min for a total of 60 minutes. The cells were washed with filtered 1% BSA in PBS pH 7.4 (2 × 1min) at room temperature, PBS pH 7.4 (2 × 1 min) at room temperature, 100 mM NaCl 50 mM Na·Bicine pH 8.3 buffer (3 × 1min) and reacted with 2.5 mM DAB (D8001-10G, Sigma-Aldrich), 4 mM H_2_O_2_ (diluted from 30% stock, H-325-100, Fisher) in 100 mM NaCl 50 mM Bicine pH 8.3. The oxidation reaction was stopped when light brown precipitate occurred after 2 to 3 minutes. The labeled cells were processed for conventional transmission electron microscopy (see below).

### Labeling and Nd-DAB2 oxidation of EdU-treated SEA cells with Cy5 azide and Fe-TAML azide

Primary small airway epithelial cells (Lonza, CC-2547) with telomerase expression (hTERT-SAEC)[21] were cultured in SAEC media on 35 mm poly-D-lysine coated MatTek dishes containing glass bottom No. 0 coverslips. SEAC were incubated with 10 μM EdU for 12 hours and then fixed with 2% glutaraldehyde in 0.1 M sodium cacodylate buffer, pH 7.4 containing 1 mM CaCl_2_ for 5 minutes at 37^°^C and then on ice for 55 minutes. Fixative was removed and cells were washed with 0.1 M sodium cacodylate buffer pH 7.4 (5 × 1 min) on ice, washed with PBS pH 7.4 (2 × 1 min) at room temperature and rinsed (2 × 1 min) with filtered 1% BSA in PBS pH 7.4 at room temperature. Click reaction of the cells was carried out at room temperature and protected from light by using a mixture of 1.0 μM Cy5 azide and 28 μM Fe-TAML azide solution freshly prepared from 900 μl click buffer (100 mM NaCl 50 mM Na·HEPES pH 7.6) 10.0 μl CuSO_4_ (100 mM in water), and the reaction initiated with 100 *μ*l of freshly prepared aqueous sodium ascorbate (100 mM) with a second 100 *μ*L aliquot added after 30 min for a total of 60 minutes. The cells were washed with filtered 1% BSA in PBS pH 7.4 (2 × 1min) at room temperature and PBS pH 7.4 (5 × 1 min) at room temperature. Fluorescence imaging of the labeled cells with Cy5 azide were collected by a Leica SPEII confocal microscope. The cells were washed with 100 mM NaCl 50 mM Na·Bicine pH 8.3 (2 × 2 min) and reacted with 2.5 mM Neodymium-DAB (Nd-DAB2), 40 mM H_2_O_2_ (diluted from 30% stock) in 50 mM Bicine 100mM NaCl pH 8.3 for 15 minutes.

### Two color TEM labeling of E4-ORF3-mSOG transduced SAEC cells with Ce-DAB2 by photooxidation and DNA labeled with Nd-DAB2 by Fe-TAML azide oxidation

Primary small airway epithelial cells (Lonza, cat. no. CC-2547) with telomerase expression (hTERT-SAEC)[21] were cultured in SAEC media on 35 mm poly-D-lysine coated MatTek dishes containing glass bottom No. 0 coverslips. Cells were infected by Adenovirus expressing E4-ORF3-mSOG[21] at multiplicity of infection (MOI) of 10. After 24 hours, infected cells were treated with 10 mM EdU for 12 hours, then fixed with 2% glutaraldehyde in 0.1 M sodium cacodylate pH 7.4 containing 1 mM CaCl_2_ for 5 minutes at 37^°^C and then on ice for 55 minutes. Fixative was removed and cells were washed with 0.1M sodium cacodylate pH 7.4 (5 × 1 min) on ice. The cells were treated with a blocking solution of 20 mM glycine (G-7126, Sigma), 10 mM potassium cyanide (31252, Sigma-Aldrich), 10 mM amino-1,2,4-trizole (8056-25G, Sigma-Aldrich) and 0.4 mM hydrogen peroxide for 30 minutes to prevent non-specific background with Ce-DAB2. The block was removed, and cells were rinsed 3 times quickly with 0.1 M sodium cacodylate pH 7.4. Cells were incubated for 30 minutes with a filtered (0.22 μm Millex 33mm PES sterile filter) 5 mM mersalyl acid solution (M9784-1G, Sigma-Aldrich) in 0.1 M sodium cacodylate pH 7.4 to block non-specific oxidation of Ce-DAB2 in mitochondria. Regions of interest were identified by standard FITC filter set (EX470/40, DM510, BA520) with a light intensity of 10% from a 150 W xenon lamp to prevent photo-bleaching and confocal fluorescence images of the adenovirus E4-ORF3-mSOG were taken by a Leica SPEII. After confocal imaging, the microscope x-y coordinates were saved. Ce-DAB2 solutions were prepared as previously described[19] and cells were pre-equilibrated for 15 minutes following filtration with a 0.22 μm Millex 33 mm PES sterile filter at room temperature. Photooxidation was performed on the previously imaged cells by blowing medical grade oxygen over the Ce-DAB2 solution and cells were illuminated at 100% intensity using the standard FITC filter set. After a light brown precipitate occurred after 6 to 7 minutes, the illumination was stopped. The Ce-DAB2 solution was removed, then cells were rinsed with 0.1 M sodium cacodylate buffer pH 7.4 (5 × 1 min) on ice, washed with PBS pH 7.4 (2 × 2 min) at room temperature and rinsed 2 × 1 min) with freshly made 2% BSA in PBS pH 7.4, at room temperature. Cells were then reacted with Fe-TAML azide and Cy5 azide using the protocol above. The cells that were previously photooxidized were then imaged by Lieca SPEII confocal microscope for Cy5 azide labeled cells. After imaging, Nd-DAB2 was precipitated by Fe-TAML catalyzed H_2_O_2_ oxidation as described in the single-color EM protocol above and reaction was stopped when light brown precipitate occurred after 15 minutes. The labeled cells were processed for conventional transmission electron microscopy, EELS, EFTEM (see below).

### CA-Fe-TAML-rhodamine labeling of Hos cells stably expressing histone H2B-HaloTag or transiently expressing HEK293T cells and oxidation of DAB

Human osteosarcoma (Hos) cells stably expressing H2B-haloTag were cultured on 35 mm glass bottom No. 0 coverslips coated with poly-d-lysine MatTek dishes in DMEM medium supplemented with 10% FBS (fetal bovine serum) without antibiotic. The H2B-HaloTag stable cells were fixed with 2% glutaraldehyde in 0.1 M sodium cacodylate pH 7.4, rinsed once with the fixative at 37^°^C, fixed for 5 minutes at room temperature and then 25 minutes on ice. After fixation, cells were rinsed with 0.1 M sodium cacodylate pH 7.4 with 0.05% glycine (5 × 2 min) at 4^°^C. Cells were treated with 0.1% saponin with 0.05% glycine in 0.1 M sodium cacodylate pH 7.4 for 20 minutes on 4^°^C and rinsed with 0.1 M sodium cacodylate pH 7.4 with 0.05% glycine (5 × 1 min) at 4^°^C. Cells were labeled with 1.0 μM CA-Fe-TAML-rhodamine in 100 mM NaCl 50 mM Na·HEPES pH 7.6 for 3 hrs at room temperature. Cells were rinsed with filtered 0.1 M sodium cacodylate pH 7.4 (3 × 1 min) containing 2% BSA at 4^°^C and 0.1 M sodium cacodylate pH 7.4 (3 × 1 min). Cells labeled with H2B-CA-Fe-TAML-rhodamine were imaged for fluorescence by a Leica SPEII confocal microscope. Cells were rinsed with 100 mM NaCl 50 mM bicine pH 8.3 (3 × 1min) at 4^°^C and incubated with 2.5 mM DAB 40mM H_2_O_2_ 100 mM NaCl 50 mM bicine pH 8.3 overnight at 4^°^C. The next day cells were then rinsed with (2 × 2 min) 100 mM NaCl 50 mM bicine pH 8.3 at 4^°^C and (3 × 2 min) 0.1M sodium cacodylate pH 7.4. The labeled cells were processed for conventional transmission electron microscopy (see below).

HEK293T cells were plated onto MatTek 35 mm glass bottom No. 0 coverslips coated with poly-d-lysine and cultured (in DMEM supplemented with 10% FBS at 5% CO_2_. On reaching 70% confluency and two days before the experiment, cells are transfected with 0.5 μg of HaloTag-H2B plasmid DNA with lipofectamine 3000 (ThermoFisher Scientific) following the manufacturer protocol. The cells were fixed using the above protocol and incubated with 1 μM CA-Fe-TAML-rhodamine in 100 mM NaCl 50 mM Na·HEPES pH 7.6 0.1% saponin for one hour at room temperature. After incubation, the cells were washed (2 × 1 min) with filtered 2% BSA in PBS, 1X pH 7.4, at room temperature and PBS, 1X pH 7.4 at room temperature. Fluorescence imaging of the cells labeled for H2B-CA-Fe-TAML-rhodamine were acquired by a Leica SPEII confocal microscope. The cells were rinsed with 100 mM NaCl 50 mM Na·Bicine pH 8.3 (2 × 2 min) and reacted with filtered 2.5 mM DAB 40 mM H_2_O_2_ 50 mM Bicine 100 mM NaCl pH 8.3 for 40-60 minutes. The labeled cells were processed for conventional transmission electron microscopy, EELS, EFTEM and electron tomography (see below).

### Labeling of microtubules using mouse anti-β-tubulin primary monoclonal antibody with secondary Fe-TAML-peg4-Cy5-goat anti-mouse IgG conjugate and oxidation of DAB with H_2_O_2_

HEK293T cells were cultured on imaging plates containing poly-D-lysine coated glass bottom No. 0 coverslips in DMEM supplemented with 10% fetal bovine serum. Cells were rinsed (x3) with cytoskeleton stabilizing buffer, CSB (10 mM Pipes buffer/150 mM NaCl/5 mM EGTA/5 mM MgCl_2_/5 mM glucose monohydrate, pH 6.8) at 37^°^C and fixed with 4% paraformaldehyde and 0.05% glutaraldehyde in CSB at 37^°^C for 5 min and then for 25 min at 4^°^C. After fixation, cells were first washed with CSB (5 × 1 min) at 4^°^C and then treated with 0.1% saponin and 0.05% glycine in CBS for 20 mins at 4^°^C while on a rocker. Cells with washed in CSB buffer with 0.05% glycine (3 × 1 min) at 4^°^C, blocked with 1% BSA, 1% normal goat serum (NGS) and 0.05% glycine in CSB for 20 min at 4^°^C and then incubated with primary mouse monoclonal antibody to *β*-tubulin (300-fold dilution, clone Tub2.1, T5201, Sigma) for 3 h at 4^°^C in 1% BSA, 1% NGS and 0.05% glycine in CSB buffer at 4^°^C. Primary antibody was removed by washing with 1% BSA 1% NGS and 0.05% glycine in CSB (5 × 3 min) at 4^°^C and then incubated with secondary Fe-TAML-peg4-Cy5-goat anti-mouse IgG conjugate in 1% BSA (0.15 ml diluted to 1ml) in 1% NGS and 0.05% glycine in CSB for overnight at 4^°^C. Cells were then washed (5 × 1 min) with CSB on ice, fixed with 2% glutaraldehyde in CSB for 20 min at 4^°^C, washed (5 × 1min) with CSB on ice and imaged on a Leica SPEII for fluorescence labeling. 5.4 mg of DAB was dissolved in 1.0 ml of 0.1 N HCl and 9.0 ml of 50 mM Bicine 100 mM NaCl pH 8.3 was added with 10 μl H_2_O_2_ (final, 40 mM from 30% stock) to the DAB solution. Cells were washed (2 × 2 min) with 50 mM Bicine 100 mM NaCl pH 8.3 on ice and the DAB/H_2_O_2_ solution was added to the cells by a 0.22μm Millex 33mm PES sterile filter (SLGSR33RS, Sigma-Aldrich) on ice. Reaction times were 60-90 min. The DAB solution was removed, and cells were washed with 50 mM Bicine 100mM NaCl pH 8.3 (2 × 2 min) on ice and 0.1 M sodium cacodylate buffer pH 7.4 (3 × 2 min). A final primary fixation with 2% glutaraldehyde in 2 mM CaCl_2_ 0.1 M sodium cacodylate pH 7.4 was done for 30 minutes on ice. The fixative was removed, and cells were washed with 0.1 M sodium cacodylate pH 7.4 (5 × 2 min) on ice.

### Post-labeling cell processing for both conventional TEM, EELS, and EFTEM

All cells after labeling experiments were posted fixed with 1% osmium tetroxide (19150, Electron Microscopy Sciences) containing 0.8% potassium ferrocyanide, 2 mM CaCl_2_ and in 0.1 M sodium cacodylate pH 7.4 for 30 minutes. Cells were washed with ddH2O at room temperature (5 × 2 min) followed by an ice-cold graded dehydration ethanol series of 20%, 50%, 70%, 90%, 100% (anhydrous) for one minute each and 2 × 100% (anhydrous) at room temperature for 1 minute each. Cells were infiltrated with one-part Durcupan ACM epoxy resin (44610, Sigma-Aldrich) to one-part anhydrous ethanol for 30 minutes, 2 times with 100% Durcupan resin for 2 hours each, a final change of Durcupan resin and immediately placed in a vacuum oven at 60^°^C for 48 hours to harden.

Cells were identified, cut out by jewel saw and mounted on dummy blocks with Krazy glue. Coverslips were removed and 70-100 nm specimen sections were created with a Leica Ultracut UCT ultramicrotome and Diatome Ultra 45^°^ 4mm wet diamond knife. Sections were picked up with 50 mesh gilder copper grids (G50, Ted Pella, Inc). DAB labeled sections were imaged by FEI Spirit transmission electron microscope at 80kV, Tietz TemCam F-224 2k by 2k CCD camera and by Serial EM software version 3.1.1a for conventional transmission electron microscopy.

The energy filtered transmission electron microscopy (EFTEM) of sections containing Ce-DAB2 and Nd-DAB2 labeling were performed by JEOL JEM equipped with an in-column Omega Filter and operating at 300kV. The samples were pre-irradiated at a low magnification of 100x for about 30 mins to stabilize the sample and minimize contamination[36]. The elemental maps for Ce and Nd were acquired at the M_4,5_ edge, the onset of which occurs at 883 eV and 978 eV, respectively[20]. For the single-color Nd-DAB2 DNA map (see Figure 2), the map was computed by the 3-window method[37], with each pre-edge and post-edge being acquired as a set of 10 individual images instead of one long exposure in order to mitigate effects of sample drift and microscope instabilities and were later aligned and summed[38]. The individual pre/post-edge image was acquired for an exposure of 120 sec, at a dose rate of ∼270 e^-^/nm^2^.sec. The total dose for the collection of the map was ∼ 9.9 × 10^5^ e^-^/nm^2^. The 2-color map of EdU-treated SEA cells transduced with E4-ORF3 (see Figure 3), was computed with the 5-window method[19], with each pre/post-edge being acquired as a set of 6 individual images for an exposure time of 180 sec. The dose rate for the acquisition was ∼940 e^-^/nm^2^.sec and the total dose for the acquisition of the map was ∼ 6.1 × 10^6^ e^-^/nm^2^. All the images and maps were acquired using an Ultrascan 4000 CCD detector from Gatan (Pleasanton, CA, USA), the maps were acquired for a hardware bin of 4 × 4 pixels. The individual images were aligned using the Template Matching plugin in ImageJ[39], and the elemental maps were computed using the EFTEM-TomoJ plugin of ImageJ[40].

## ACKNOWLEDGEMENTS

We thank Hiroyuki Hakozaki for technical assistance, Horng Ou and Clodagh O’Shea (Salk Institute) for providing E4-ORF3-mSOG transduced SAEC cells, and Luke Lavis (Janelia, HHMI) for the kind gift of HaloTag protein.

## AUTHOR CONTRIBUTIONS

S.R.A., J.N., D.B, and M.H.E. designed research; M.M., R.R., G.A.C., S.P., T.J.D., J.H., and S.R.A. performed research; M.M., R.R., G.A.C., S.R.A., D.B., S.P., J.N., and M.H.E analyzed data; J.N., S.R.A., and M.H.E. drafted the paper and all authors revised it.

---

**Figure S1.**
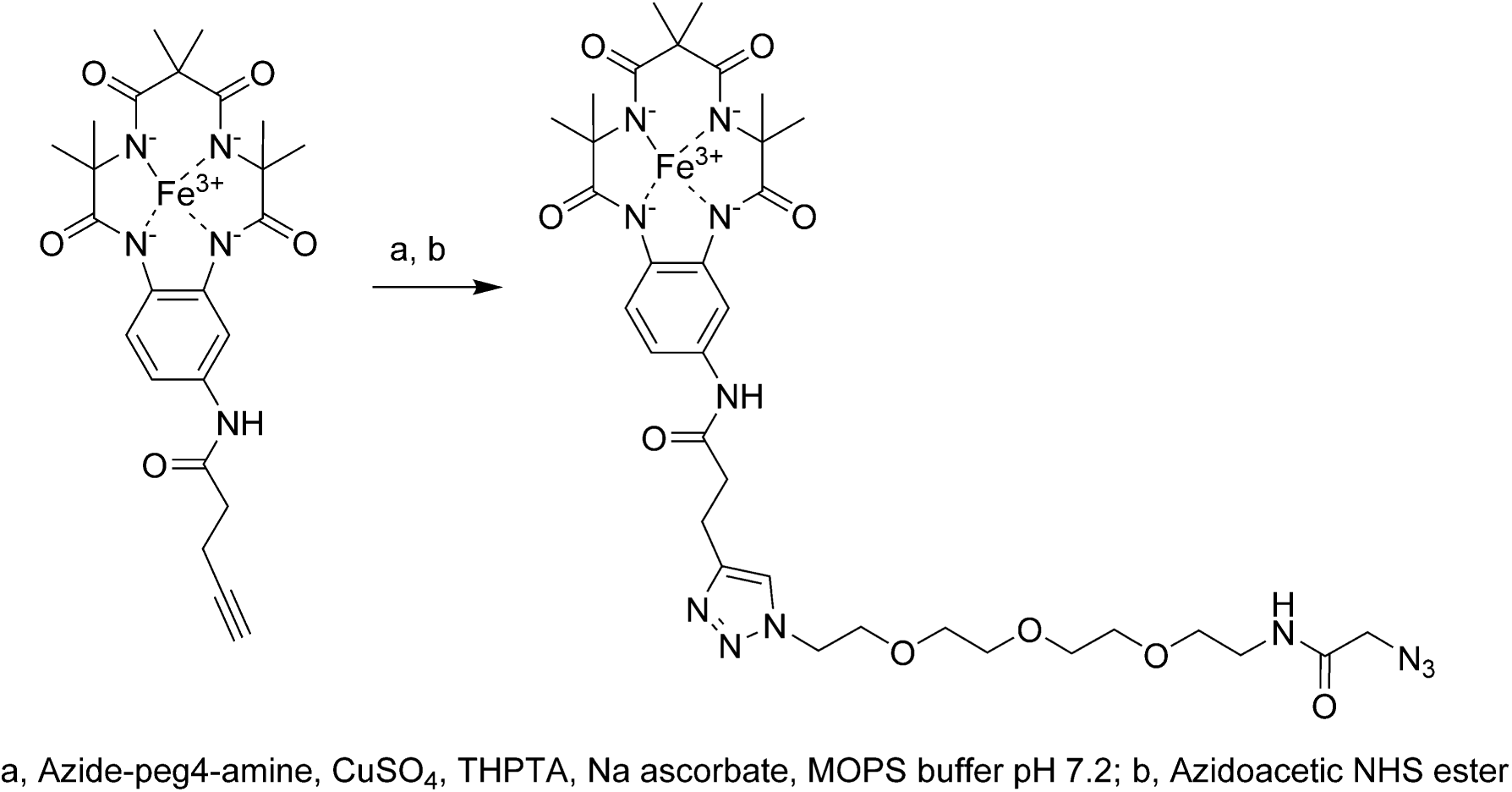
Synthesis of Fe-TAML azide.

**Figure S2.**
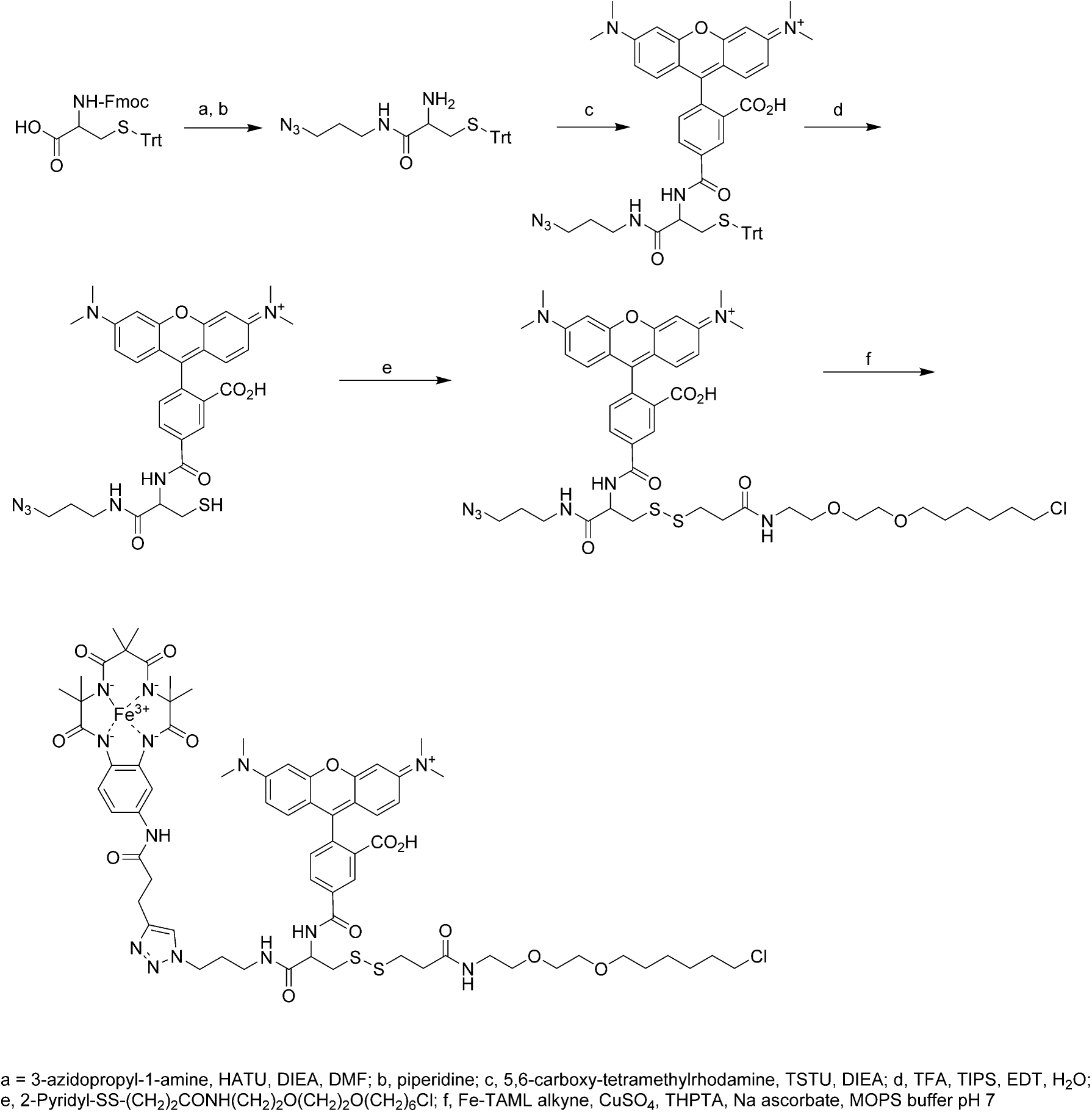
Synthesis of CA-Fe-TAML-rhodamine.

**Figure S3.**
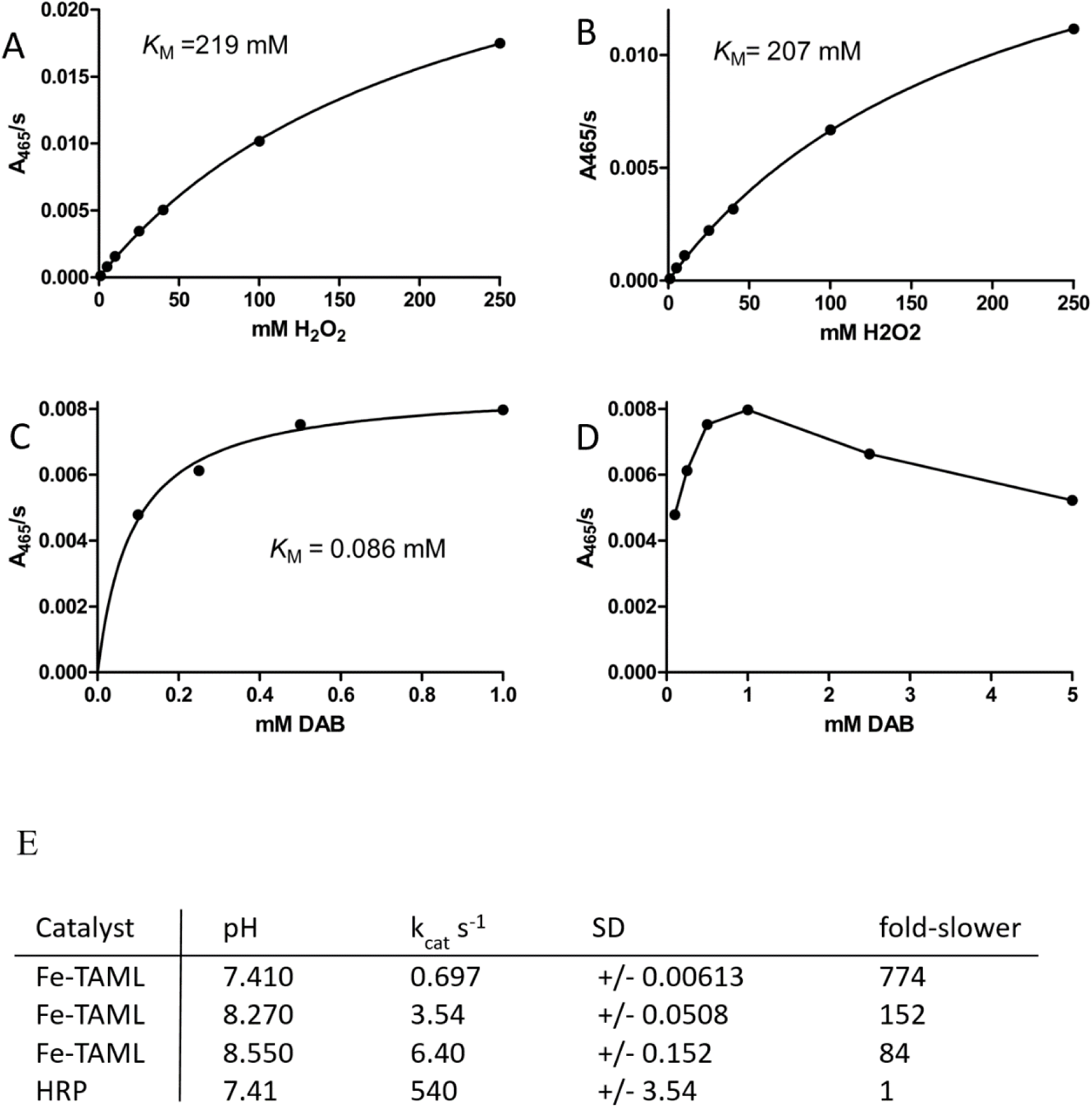
Fe-TAML kinetics. Determination of *K*_M_ values for H_2_O_2_ and DAB at pH 7.4 and 8.3. A) H_2_O_2_, pH 7.4. B) H_2_O_2_, pH 8.3. C) DAB, pH 7.4. D) Rates at higher DAB concentrations. E) Rate constants and relative rates of Fe-TAML B* and HRP derived from Figure 1H.

**Figure S4.**
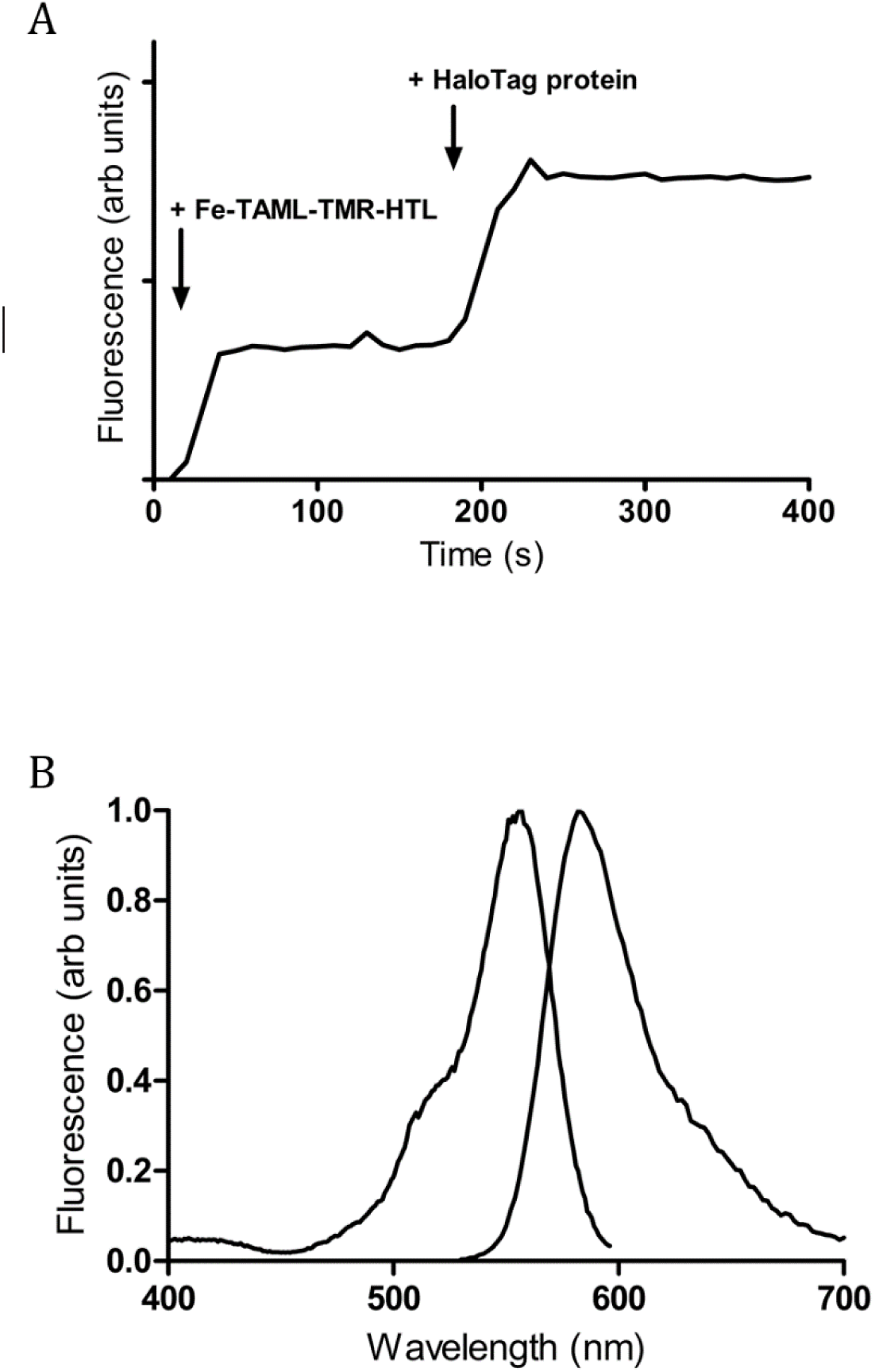
A) Fluorescence time course for reaction of Fe-TAML-TMR-HTL with HaloTag protein in vitro. B) Fluorescence spectra of resulting conjugate

